# Human FcγRIIIa activation on splenic macrophages drives the in vivo pathogenesis of dengue disease

**DOI:** 10.1101/2022.11.02.514909

**Authors:** Rachel Yamin, Kevin S. Kao, Margaret R. MacDonald, Tineke Cantaert, Charles M. Rice, Jeffrey V. Ravetch, Stylianos Bournazos

## Abstract

Although dengue virus (DENV) infection typically causes asymptomatic disease, DENV-infected patients can experience severe complications. A risk factor for symptomatic disease is pre-existing anti-DENV IgG antibodies. Cellular assays suggested that these antibodies can enhance viral infection of Fcγ receptor (FcγR)-expressing myeloid cells. Recent studies, however, revealed more complex interactions between anti-DENV antibodies and specific FcγRs by demonstrating that modulation of the IgG Fc glycan correlates with disease severity. To investigate the *in vivo* mechanisms of antibody-mediated dengue pathogenesis, we developed a mouse model for dengue disease that recapitulates the unique complexity of human FcγRs. Our studies reveal that the *in vivo* pathogenic activity of anti-DENV IgG antibodies is exclusively mediated through engagement of FcγRIIIa expressed on splenic macrophages, resulting in inflammatory sequelae and mortality. These findings highlight the importance of IgG-FcγRIIIa interactions in dengue disease, with important implications in the design of safer vaccination approaches and effective therapeutic strategies.

---

Dengue virus (DENV) poses a significant threat to global public health, causing an estimated 50 to 100 million new infections annually^1^. Although DENV infection often manifests as an inapparent or mildly symptomatic disease, approximately 10% of DENV-infected patients experience more severe forms of symptomatic disease, which include dengue hemorrhagic fever (DHF) and dengue shock syndrome (DSS), characterized by persistent high-grade fever, thrombocytopenia, and vascular leakage, resulting in significant morbidity and mortality^2,3^.

Several genetic and immunological determinants have been proposed as key susceptibility factors to symptomatic dengue disease^4-6^. Large scale epidemiological studies in DENV-endemic areas have previously revealed a higher prevalence of symptomatic dengue disease in patients with prior history of DENV infection, as well as in newborns from DENV-immune mothers^7,8^. These observations have been interpreted to suggest that dengue disease pathogenesis is primarily driven by circulating, pre-existing serum IgG antibodies against DENV, which are elicited in response to prior DENV infection or transferred from DENV-immune mothers to neonates.

Although a pathogenic role for anti-DENV antibodies is supported by substantial epidemiological evidence, the mechanisms by which these antibodies contribute to dengue disease susceptibility have not been well characterized. It has been previously hypothesized that pre-existing antibodies that bind but do not neutralize the DENV virions generate immune complexes that are recognized by Fcγ receptor (FcγR)-expressing leukocytes, leading to increased virus infectivity^6^. This phenomenon, termed antibody-dependent enhancement (ADE), has been demonstrated *in vitro* using FcγR-expressing cell lines, such as K562 and U937^9-13^. While at high antibody concentrations infection is blocked due to antibody-mediated neutralization, at sub-neutralizing levels, anti-DENV antibodies promote DENV infection, suggesting that the titers of anti-DENV antibodies likely modulate ADE of dengue infection. Consistent with this hypothesis, epidemiological studies revealed that the titers of anti-DENV antibodies are associated with dengue disease susceptibility, as higher prevalence of symptomatic disease has been observed in patients with sub-neutralizing anti-DENV titers^14,15^.

Despite such evidence, anti-DENV antibody titers alone cannot explain why among symptomatic patients there is such a wide spectrum of disease severity, which ranges from the mild febrile illness seen in classical dengue fever (DF) cases to the life-threatening DSS^6^. Therefore, additional determinants likely exist that modulate the pathogenic activity of anti-DENV antibodies, including the Fc domain heterogeneity of these antibodies and FcγR expression on specific cellular subsets, which collectively impact their capacity to engage and activate FcγRs. Five canonical FcγRs are expressed on human immune cells, FcγRI, FcγRIIa, FcγRIIb, FcγRIIIa, and FcγRIIIb and are regulated in response to specific cytokine stimulation. Differential engagement of specific FcγRs is determined by both the IgG subclass and Fc glycan composition and will dictate whether immune complex engagement results in pro-or anti-inflammatory responses^16,17^. Indeed, in two independent cohorts, it has been demonstrated that IgG antibodies from symptomatic dengue disease patients are characterized by elevated levels of afucosylated Fc glycoforms, which exhibit higher affinity for FcγRIII^18,19^. Importantly, the abundance of afucosylated IgG antibodies was not only associated with symptomatic disease, but also correlated with disease severity, as DF and DSS patients had the lowest and the highest levels of afucosylated Fc glycoforms, respectively^19^.

Although these findings support an association between afucosylated IgG antibodies and dengue disease susceptibility and severity, definitive evidence on the contribution of these antibody glycoforms to the *in vivo* pathogenesis of dengue disease has not been demonstrated. Likewise, the role of specific FcγR pathways during dengue disease remains unknown, mainly due to the lack of biologically relevant experimental model systems that recapitulate the unique complexity of human FcγRs. Indeed, substantial interspecies differences in the affinity, structure, function, and expression pattern of FcγRs between humans and animal species that are commonly used in biomedical research, represent a major obstacle in efforts to understand the *in vivo* biological activity of human IgG antibodies^20,21^.

To overcome this limitation and study the role of human FcγRs during dengue disease pathogenesis, we developed an *in vivo* model of dengue disease that expresses the full array of human FcγRs and is permissive for DENV infection. Using this model and a panel of Fc domain engineered anti-DENV antibodies with defined affinities for specific human FcγRs, we assessed the mechanisms by which human FcγRs contribute to the antibody-mediated dengue disease. We observed a critical role for anti-DENV antibodies engineered for enhanced binding to the activating receptor FcγRIII in driving disease pathogenesis, consistent with the clinical correlation of afucosylated IgG with disease severity in dengue infected patients^18,19^. In a series of cellular depletion studies of the various FcγRIII-expressing leukocyte populations, we identified splenic macrophages as a key effector cell type during dengue disease and demonstrate that their engagement results in inflammatory cytokine responses and disease sequelae.

## Results

### Comparison of in vitro models of ADE of dengue infection to assess the role of human FcγRs

Myeloid cell lines derived from leukemia patients, such as U937 and K562, have been historically used to study ADE of DENV infection and assess the requirements for FcγR engagement in this process. These cells are resistant to dengue infection in the absence of antibodies and require FcγR engagement to mediate virus uptake and subsequent replication in these cells^9,10^. Consistent with previous studies that established a key role for Fc-FcγR interactions in DENV infection of these cells, the anti-DENV monoclonal antibody (mAb) (clone C10) promoted infection of K562 (Fig. 1a) and U937 (Fig. 1b) cells, when expressed as human IgG1 Fc, but not when its Fc is modified to abrogate human FcγR binding (Fc null). However, as described above, FcγRs are a complex family of surface glycoproteins and since these cell lines do not express the full array of human FcγRs (Fig. 1c and Extended Data Fig. 1), they provide limited information on the role of specific human FcγRs during ADE of DENV infection. For example, the lack of FcγRIIIa expression on these cells results in their inability to detect enhanced ADE for afucosylated glycoforms (Fig. 1a,b), the IgG species which have been previously reported to be associated with dengue disease severity^18,19^. To address this limitation, we engineered the U937 cell line (U937-panFcγR) to express all classes of human FcγRs (Fig. 1c) and compared the *in vitro* ADE activity between fucosylated and afucosylated Fc glycoforms of anti-DENV mAbs. Afucosylated Fc glycoforms induce enhanced infection of U937-panFcγR compared to their fucosylated counterparts (Fig. 1d), supporting a role for FcγRIIIa in mediating ADE of DENV infection *in vitro*.

**Fig. 1:**
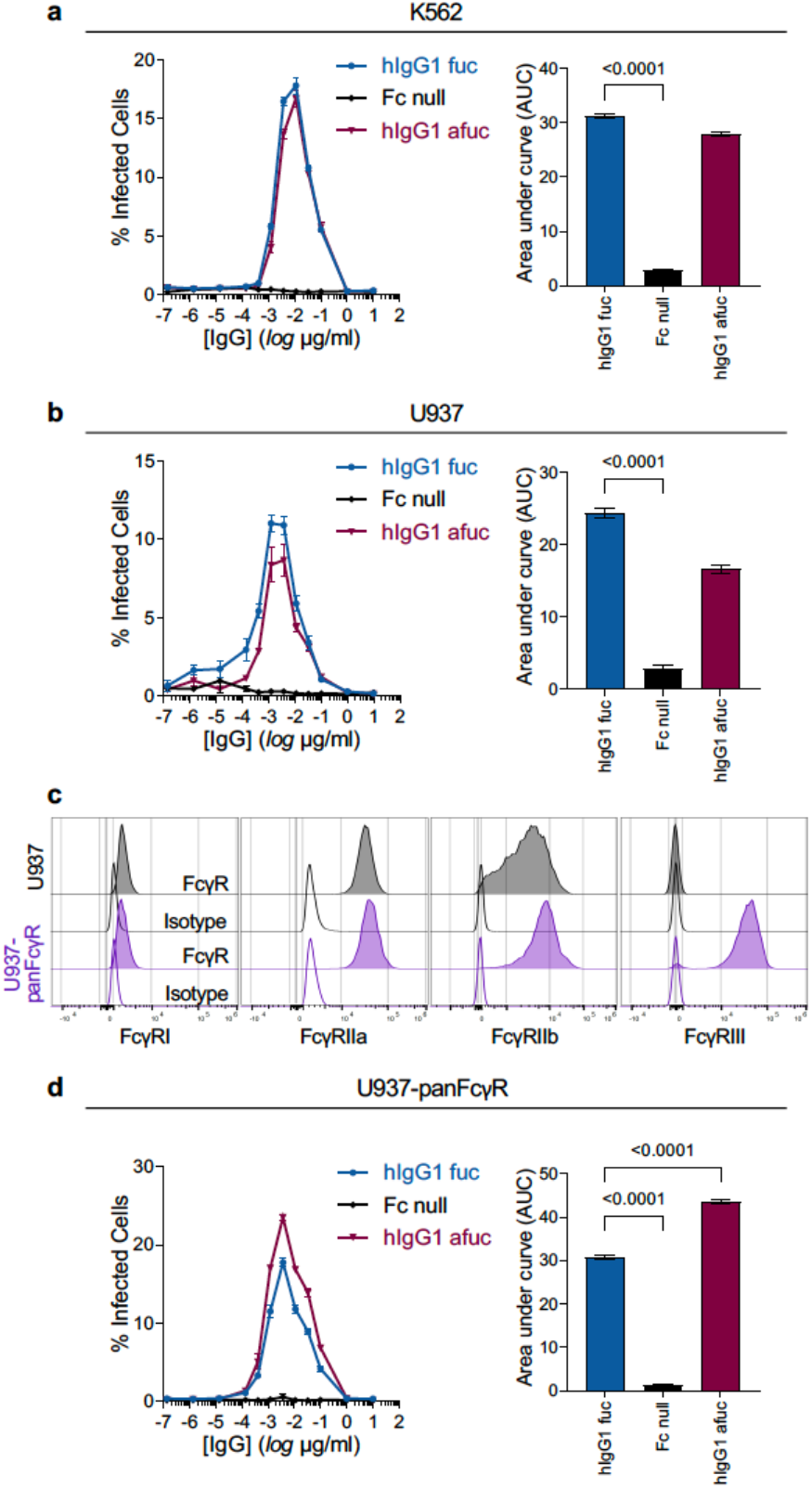
*In vitro* ADE activity of anti-DENV IgG antibodies is dependent on the FcγR expression profile of the cell lines used in ADE assays. The *in vitro* ADE activity of the anti-DENV mAb C10 expressed as human IgG1 fucosylated (fuc) or afucosylated (afuc), or as Fc variant with diminished binding to all classes of human FcγR (Fc null) was assessed in K562 (**a**), U937 (**b**), and U937-panFcγR (**d**) cells. Percent (%) infection was assessed by flow cytometry. Area under curve (AUC) was calculated for each variant and compared to fucosylated hIgG1 by one-way ANOVA (Bonferroni post hoc analysis adjusted for multiple comparisons). Figure shows one representative experiment out of two performed in triplicates. (**c**) FcγR expression on U937 (gray) and U937-panFcγR (purple) was assessed by flow cytometry. See also Extended Data Fig. 1 for FcγR expression profile of K562 cells.

### Development of Ifnar1^-/-^ mice humanized for all classes of FcγRs for the study of human FcγR pathways during ADE of dengue disease

*In vitro* ADE experimental systems only assess the capacity of anti-DENV antibodies to mediate the uptake of DENV virions by FcγR-expressing cells, thereby leading to increased infectivity and viral replication. However, antibody-mediated enhancement of viral replication alone cannot account for the complex pathophysiological features of symptomatic dengue disease. Therefore, understanding of the mechanisms by which anti-DENV IgG antibodies mediate pathogenic activities necessitates the use of *in vivo* models that mirror the clinical features of symptomatic dengue disease.

Since wild-type mice are resistant to dengue infection, the current *in vivo* models of dengue disease rely on the disruption of type I interferon signaling (IFNα/βR-KO) to permit dengue replication and pathogenesis^22^. Such mouse strains develop symptomatic dengue disease upon DENV challenge and have been historically used for the study of DENV infection *in vivo*. While these strains provide a significant resource to investigate many aspects of dengue disease pathogenesis, they are inadequate for the investigation of the role of pre-existing antibodies in the disease enhancement seen in symptomatic disease. The major differences in the expression pattern, structure, and diversity between mouse and human FcγRs represent significant barriers to the interpretation of how anti-dengue antibodies elicited in response to dengue infection or vaccination in human cohorts contribute to disease severity through human FcγR engagement. To overcome these limitations, we generated IFNα/βR-KO (*Ifnar1*^-/-^) mouse strains on a background of either FcγR deficient (FcγR null) or FcγR humanized mice, the latter of which generally recapitulate the expression pattern of human FcγRs found on human immune cell populations^20^. Characterization of these mouse strains verified the deletion of the *Ifnar1* gene and confirmed the appropriate expression of human FcγRs on the various effector leukocyte populations (Extended Data Fig. 2).

To validate this new mouse model and determine its utility for studying ADE in dengue infection and disease, we administered polyclonal IgG samples derived from symptomatic dengue patients^19^ to *Ifnar1*^-/-^/FcγR humanized mice or *Ifnar1*^-/-^/FcγR KO mice prior to DENV infection. This experimental scheme was selected to mimic the presence of pre-existing IgG antibodies seen in secondary DENV infection cases (Fig. 2a). *Ifnar1*^-/-^/FcγR KO mice were fully protected from DENV disease, whereas *Ifnar1*^-/-^/FcγR humanized mice showed evidence of disease, succumbing to death a few days post-DENV infection (Fig. 2b,c). Additionally, antibody-treated *Ifnar1*^-/-^/FcγR humanized mice develop severe thrombocytopenia, a clinical feature that is characteristic of severe dengue disease in humans (Fig. 2d). Histological evaluation of various organs from anti-DENV-treated *Ifnar1*^-/-^/FcγR humanized mice revealed massive hepatocellular necrosis, extensive neutrophilic inflammation in skin, lymph nodes, and intestine, as well as thrombosis in multiple organs, such as the intestine and the lungs (Fig. 2e). Such histopathological findings are consistent with those observed in postmortem evaluations of dengue-related deaths in humans^23,24^.

**Fig. 2:**
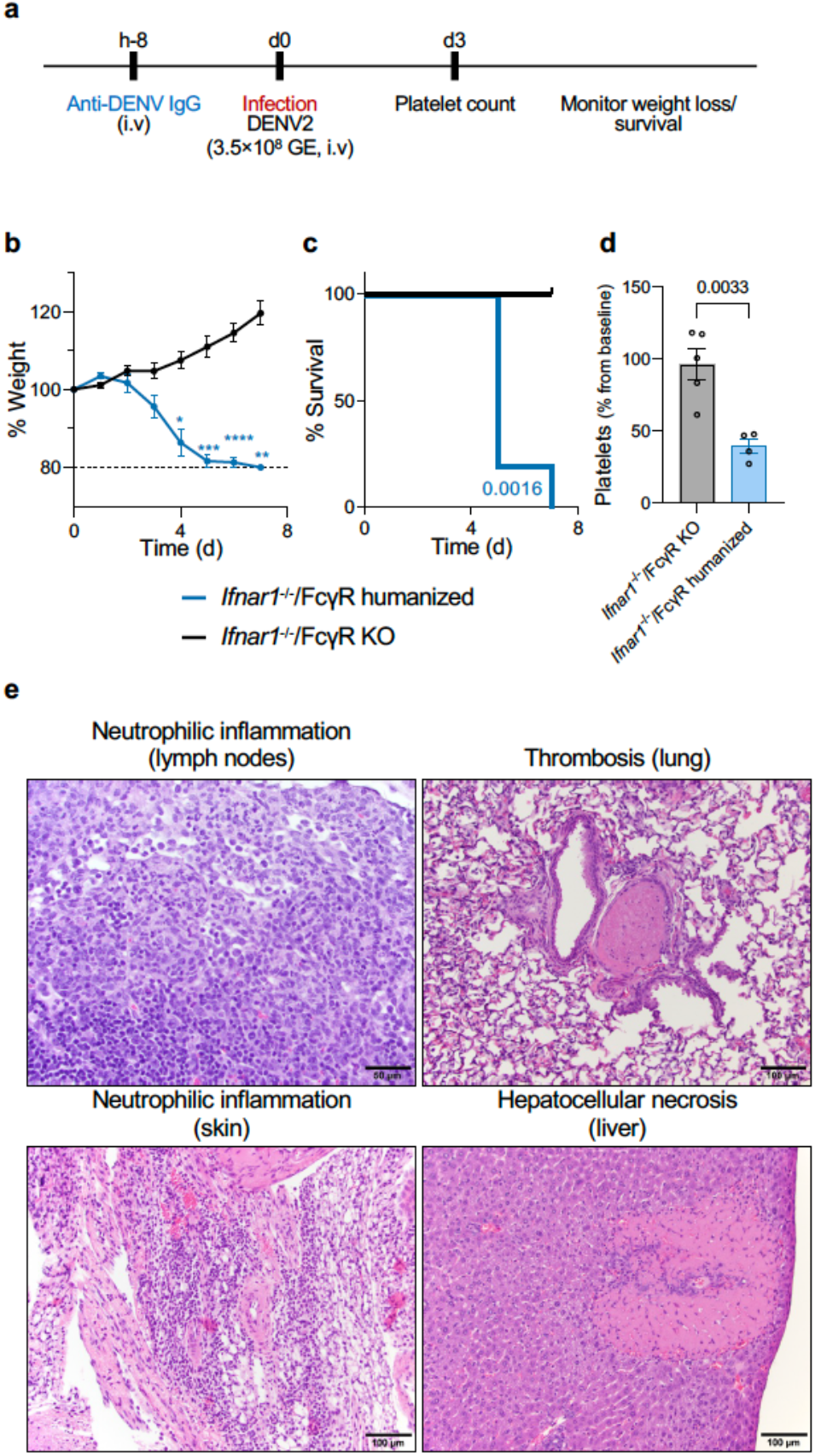
Development of experimental models for the study of human FcγR pathways during ADE of dengue disease. (**a**) Overview of the experimental strategy to assess *in vivo* ADE of dengue disease in *Ifnar1*^-/-^ /FcγR KO or *Ifnar1*^-/-^/FcγR humanized mice. (**b**,**c**) Following the experimental strategy in (A), *Ifnar1*^-/-^/FcγR KO (black) and *Ifnar1*^-/-^/FcγR humanized (blue) mice were treated with 200 μg of serum polyclonal human IgGs purified from DENV patients. Weight loss (**b**) and survival (**c**) were compared between the mouse lines by two-way ANOVA (Bonferroni post hoc analysis adjusted for multiple comparisons) or log-rank (Mantel-Cox) test, respectively. \**P*=0.01; \*\**P*=0.002; \*\*\**P*=0.0006; \*\*\*\**P*<0.0001 (**d**) Platelet counts were determined on day 3 post-infection and calculated as percentage from baseline (day 0). Platelet counts were compared by two-tailed unpaired t-test. (**b-d**) n = 4-5 mice/group. (**e**) Pathological changes related to DENV infection were assessed on day 4 post DENV infection following H&E staining and histological evaluation. Mice that were pre-treated with anti-DENV antibodies exhibited neutrophilic inflammation (upper left panel, 40x; lower left panel, 20x), thrombosis (upper right panel, 20x), and hepatocellular necrosis (lower right panel, 20x). Images are representative of three infected mice. See also Extended Data Fig. 2 for further characterization of *Ifnar1*^-/-^/FcγR humanized mice.

### Development of a panel of Fc domain variants for anti-DENV antibodies reveals a critical role for FcγRIII in mediating ADE of dengue infection in vitro

Humans express several FcγRs that are broadly divided into two classes: (i) activating, which contain an ITAM motif and mediate pro-inflammatory activities, and (ii) inhibitory, which through their ITIM domain limit the activity of activating FcγRs. The biological outcome of IgG-mediated signaling is determined by the affinity of the IgG Fc domain for the activating and inhibitory FcγRs co-expressed on the surface of effector leukocytes^25^. We have previously developed a panel of engineered Fc domain variants of human IgG1 that exhibit selectively enhanced affinity for specific human FcγRs^26^ (Fig. 3a and Extended Data Table 1). To assess the contribution of the various human FcγR classes to the pathogenic activity of anti-DENV IgG antibodies, we generated Fc variants for two anti-DENV mAbs: clone C10, which is reactive against all DENV serotypes^27^ and clone 2D22, which is specific for the DENV2 serotype^28^. We verified that none of these Fc modifications affect their *in vitro* neutralization activity (Fig. 3b) and assessed their capacity to mediate ADE of DENV infection using U937-panFcγR cells. Fc variants with enhanced affinity for FcγRIII (GAALIE or ALIE variants) promoted increased DENV infection *in vitro*, whereas mAbs with increased affinity for FcγRIIa (GA variant) show no enhancement over human IgG1 (Fig. 3c-e). Consistent with previous reports on the requirement for activating FcγR engagement for ADE of DENV infection *in vitro*^29^, Fc variants with enhanced affinity to the inhibitory receptor FcγRIIb (V11) or diminished affinity for all FcγR classes (GRLR) exhibit reduced or no ADE activity, respectively.

**Fig. 3:**
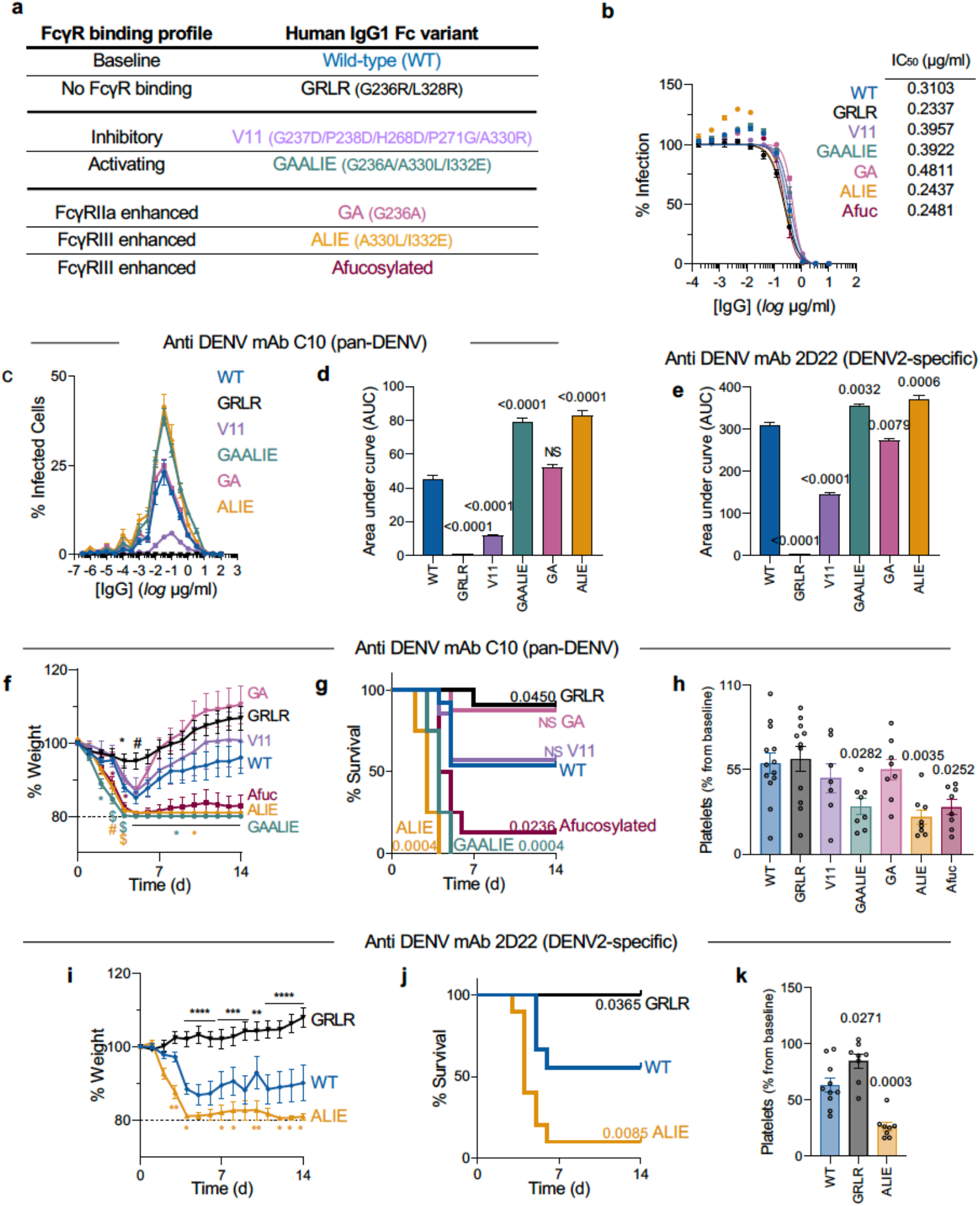
FcγRIII plays a critical role in mediating ADE of dengue infection *in vitro* and in enhanced antibody-mediated dengue disease *in vivo*. (**a**) Anti-DENV mAbs were expressed as human IgG1 Fc variants with differential affinity for specific classes of human FcγRs. See also Table S1. (**b**) Neutralization activity of C10 Fc variants was evaluated in U937-DC-SIGN cells. Infection percentage was determined 24 hours post-infection by flow cytometry. IC_50_ values were calculated from the neutralization curves for each Fc variant. n = 1 experiment performed in duplicates. *In vitro* ADE activity of C10 (**c**,**d**) or 2D22 (**e**) Fc variants was assessed in U937-panFcγR cells. Infection percentage (**c**) was determined by flow cytometry, and AUC (**d**,**e**) was calculated for each Fc variant and compared to wild-type (WT) human IgG1 by one-way ANOVA (Bonferroni post hoc analysis adjusted for multiple comparisons). Figure shows one representative experiment out of two performed in duplicates. (**f-k**) *Ifnar1*^-/-^/FcγR humanized mice were treated with 20 μg (i.v.) of Fc variants of anti-DENV mAb (F-H, C10; I-K, 2D22) according to the experimental scheme in Fig. 2a. Weight loss (**f, i**), survival (**g, j**) and platelet counts on 3 dpi (**h, k**) were compared to WT human IgG1 by two-way ANOVA (Bonferroni post hoc analysis adjusted for multiple comparisons), log-rank (Mantel-Cox) test and one-way ANOVA (Bonferroni post hoc analysis adjusted for multiple comparisons), respectively. **f**: \**P*<0.03; #*P*<0.008; $*P*<0.0001. **i**: \**P*<0.04; \*\**P*<0.004; \*\*\**P*<0.0005. \*\*\*\**P*<0.0001. See also Extended Data Fig. 3. (**f-h**) n = 7 (V11), n = 8 (GA, ALIE, GAALIE, Afuc), n = 11 (GRLR), n = 13 (WT) mice per group from two independent experiments. (I-K) n = 8 (GRLR), n = 9 (WT), n = 10 (ALIE) mice per group from two independent experiments.

### Engagement of the activating FcγRIII is associated with enhanced antibody-mediated dengue disease in vivo

Comparative analysis of the *in vitro* ADE activity of anti-DENV mAb Fc variants revealed that enhanced engagement of FcγRIII is associated with increased DENV infectivity. To determine the contribution of human FcγRs to the *in vivo* dengue disease pathogenesis, we followed the experimental scheme outlined in Fig. 2a and assessed the pathogenic activity of Fc variants of anti-DENV mAbs (C10 or 2D22) in *Ifnar1*^-/-^/FcγR humanized mice. To mimic human secondary dengue cases, antibodies were administered prior to DENV infection at doses sufficient to achieve sub-neutralizing levels (determined in titration studies; Extended Data Fig. 3). In agreement with our *in vitro* observations, wild-type human IgG1 causes enhanced disease, whereas the Fc null variant GRLR shows minimal pathogenic activity (Fig. 3f-3h), thereby confirming the requirements of Fc-FcγR interactions for dengue disease pathogenesis. Both the FcγRIIa-(GA) and the FcγRIIb-enhanced (V11) variants exhibit pathogenic activity comparable to that of wild-type human IgG1, as seen by weight loss between days 1 and 5 (Fig. 3f), survival rates (Fig. 3g), and platelet counts (Fig. 3h), suggesting a limited role for these receptors in mediating ADE of dengue disease. By contrast, when anti-DENV mAbs are expressed as Fc variants with improved affinity for FcγRIII (GAALIE, ALIE, or afucosylated), enhanced disease is observed, as evidenced by accelerated weight loss, higher mortality rate, as well as significant reduction in platelet counts. As observed for the pan-DENV mAb C10, Fc variants of the DENV2-specific mAb 2D22 enhanced for FcγRIII (ALIE) also induce more severe disease compared to the wild-type human IgG1 (baseline FcγR binding) or the GRLR (no FcγR binding) variants (Fig. 3i-k).

### FcγRIIIa^+^ splenic macrophages are necessary for ADE of dengue disease in vivo

Our findings support a role for FcγRIII in modulating the *in vivo* antibody-dependent pathogenesis of dengue disease. Since FcγRIII is expressed on a variety of innate immune cells, including NK cells, neutrophils, monocytes, and macrophages (Extended Data Fig. 2), we performed a series of cellular depletion studies in *Ifnar1*^-/-^/FcγR humanized mice to assess the contribution of these effector leukocytes to the ADE of dengue disease. Depletion of NK cells (Fig. 4a,b,k), neutrophils (Fig. 4c,d,k), or CCR2^+^ monocytes (Fig. 4e,f,k) had no impact on the pathogenic activity of FcγRIII-enhanced anti-DENV mAb Fc variants. In contrast, depletion of macrophages via intravenous (i.v.) administration of clodronate liposomes abrogated the *in vivo* pathogenic activity of FcγRIII-enhanced variants, suggesting a key role for FcγRIIIa-expressing, clodronate-sensitive macrophages during dengue disease pathogenesis (Fig. 4g,h,k).

**Fig. 4:**
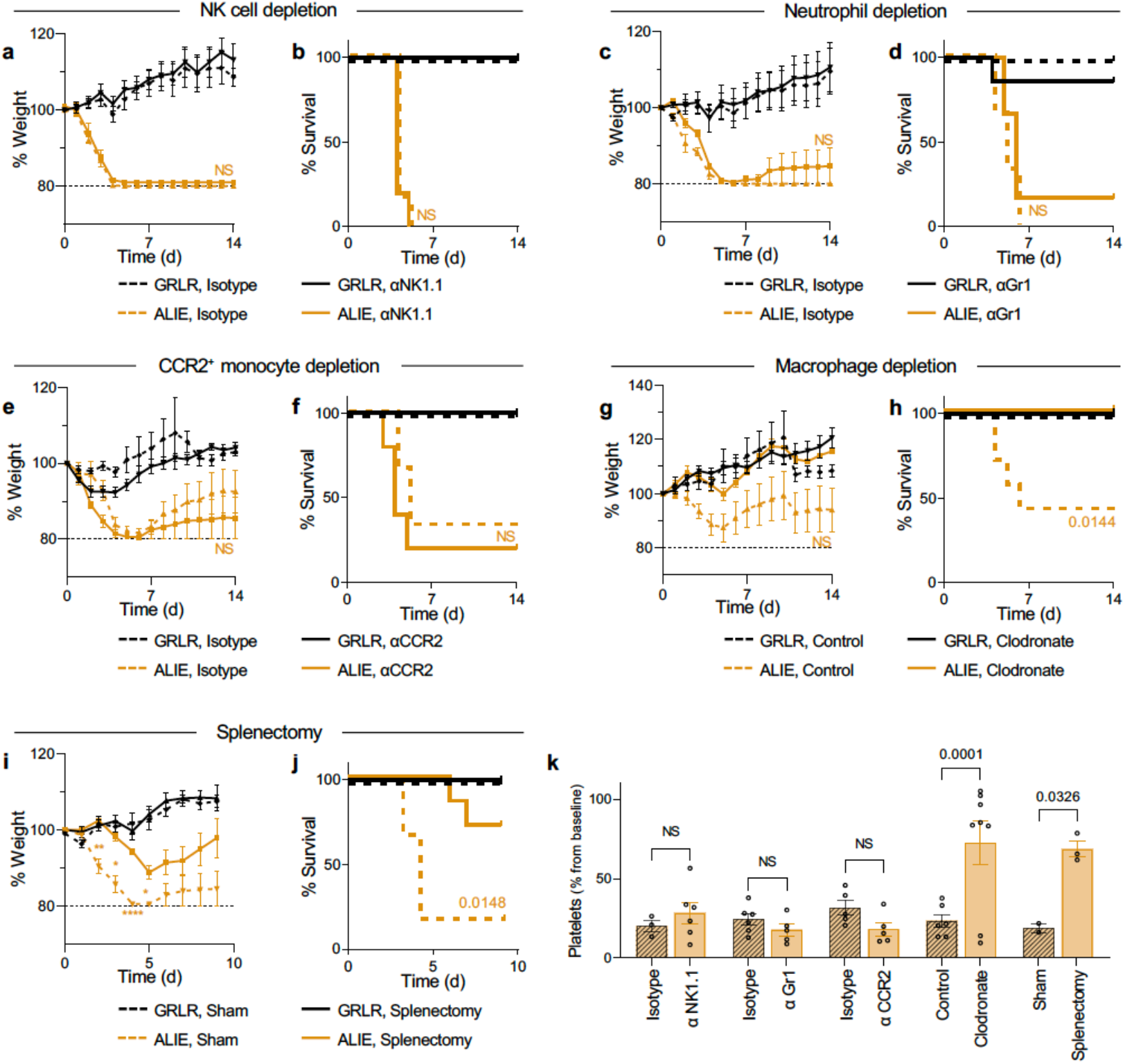
*In vivo* ADE of dengue disease is mediated by FcγRIIIa^+^ splenic macrophages. (**a-f**) *Ifnar1*^-/-^/FcγR humanized mice were administered with anti-mouse NK1.1 (**a**,**b**), anti-mouse Gr-1 (**c**,**d**) or anti-mouse CCR2 (**e**,**f**) antibodies to deplete NK cells, neutrophils, or CCR2^+^ monocytes (solid lines), respectively. Matched isotype control for each antibody is indicated in a dotted line. One day later, mice were treated with GRLR (black) or ALIE (yellow) variants of 2D22 (20 μg), followed by DENV challenge. (**g**,**h**) Macrophage depletion was achieved through administration of clodronate (solid line) or matched liposome control (dotted line) one day prior to 2D22 treatment and DENV infection. (**i**,**j**) *Ifnar1*^-/-^/FcγR humanized mice were splenectomized (solid line) and allowed to recover for 5 days prior to treatment with GRLR (black) or ALIE (yellow) variants of 2D22 and DENV infection. Weight loss and survival were compared between the ALIE groups (with or without depletion) by two-way ANOVA (Bonferroni post hoc analysis adjusted for multiple comparisons) or log-rank (Mantel-Cox) test, respectively. **i**: \**P*<0.02; \*\**P*<0.004; \*\*\*\**P*<0.0001. (**k**) Platelet counts on day 3 post-infection were determined and compared between ALIE treated mice with or without the relevant depletion. NS, not significant.

Intravenous administration of clodronate liposomes preferentially targets liver- and spleen-resident macrophages, while it has minimal impact on macrophages present in other tissues (e.g. lung)^30^. To evaluate the role of splenic macrophages, we assessed the *in vivo* pathogenic activity of FcγRIII-enhanced anti-DENV mAbs in splenectomized *Ifnar1*^-/-^/FcγR humanized mice. While administration of FcγRIII-enhanced mAbs induces lethality in sham-treated mice, we observed that the pathogenic effect of these mAbs is reduced in splenectomized mice, suggesting that the *in vivo* disease-enhancing activity of anti-DENV mAbs is likely mediated by spleen-resident macrophages (Fig. 4i-k).

### Activation of FcγRIII on splenic macrophages results in inflammatory cytokine production

To gain further insights into the mechanisms by which engagement of FcγRIIIa on splenic macrophages results in enhanced dengue disease *in vivo*, we analyzed viral titers in the spleens of DENV-infected *Ifnar1*^-/-^/FcγR humanized mice. Anti-DENV mAb Fc variant enhanced for FcγRIII binding (ALIE) promote increased DENV replication, as evidenced by the significantly higher viral titers observed in ALIE-treated mice compared to their GRLR-treated counterparts (Fig. 5a). Consistent with the higher viral titers in the spleens of ALIE-treated mice, we also observed increased frequency of DENV-infected splenic macrophages (DENV^+^/CD11b^+^/F4/80^+^) in mice that have received FcγRIII-enhanced variants (Fig. 5b,c).

**Fig. 5:**
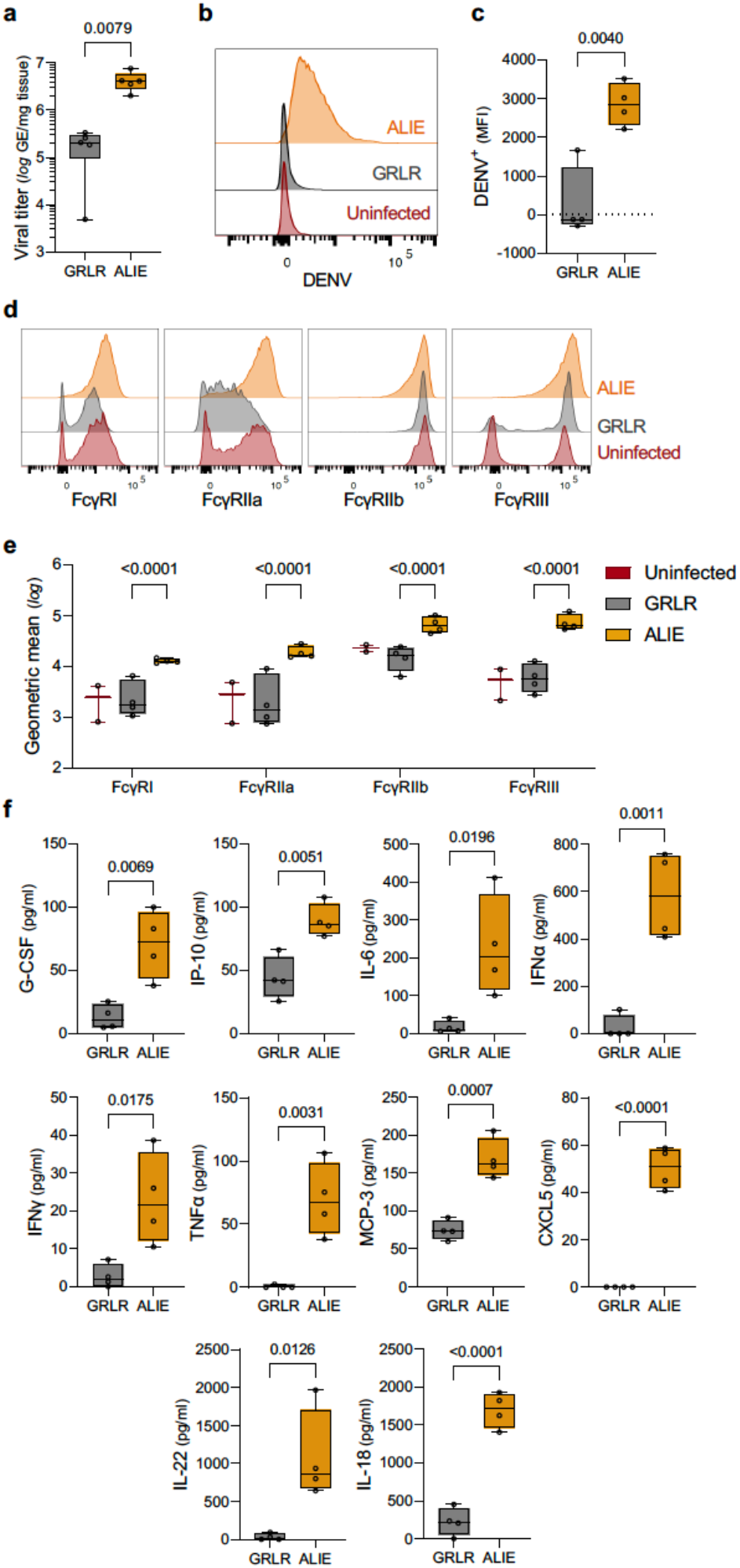
Engagement of FcγRIIIa is associated with elevated levels of inflammatory cytokines. (**a**) *Ifnar1*^-/-^/FcγR humanized mice were treated with GRLR (gray) or ALIE (yellow) variants of 2D22, followed by DENV infection. Tissue viral titers were assessed on day 3 post-infection and compared between treatment groups by one-way ANOVA (Bonferroni post hoc analysis, adjusted for multiple comparisons). DENV infection (**b**,**c**) and FcγR expression (**d**,**e**) on macrophages (CD11b^+^/F4/80^+^) in the spleen of antibody-treated DENV-infected mice were assessed on day 4 post-infection. (**b**,**d**) Representative histogram overlay plots of DENV-positive macrophages (**b**) and FcγR expression (**d**) on macrophages in the spleen. (**c**,**e**) Quantitation of DENV infection (DENV^+^; MFI, median fluorescence intensity) or FcγR expression (log geometric mean) in splenic macrophages. The respective uninfected (for DENV infection) or isotype control (for FcγR expression) were subtracted from GRLR and ALIE treated groups. Results are from 4 mice per group for GRLR and ALIE treated mice and 2 mice per group for the uninfected mice. (**f**) Serum levels of pro-inflammatory cytokines and chemokines in mice following mAb treatment and DENV infection. Levels were determined on day 3 post infection. n = 4 mice per group from one experiment performed in duplicates.

Analysis of the FcγR expression profile of splenic macrophages from *Ifnar1*^-/-^/FcγR humanized mice revealed high FcγRIIIa expression at baseline (Fig. 5d), suggesting that this macrophage population is likely the primary target for DENV infection in the presence of FcγRIII-enhanced anti-DENV mAbs. Interestingly, engagement of FcγRIII was associated with upregulation of FcγRIII expression on splenic macrophages (Fig. 5d,e). These findings suggest that activation of the FcγRIIIa pathway by ALIE variants not only results in increased infection of splenic macrophages, but also induces the activation of additional pro-inflammatory pathways that modulate leukocyte function, thereby further contributing to dengue disease pathogenesis.

Indeed, FcγRIIIa engagement was associated with a significant increase in the serum levels of several pro-inflammatory cytokines and chemokines, including TNFα, IL-18, IFNα, and IFNγ, which have also been reported to be elevated in patients with severe dengue disease (Fig. 5f)^31-33^. Given the important role of these cytokines and chemokines in modulating leukocyte differentiation and recruitment, antiviral responses, as well as macrophage activation, engagement of FcγRIII on splenic macrophages likely precipitates a dysregulated inflammatory response, characterized by excessive or inappropriate activation of multiple pro-inflammatory cascades that drive dengue disease pathogenesis.

## Discussion

A characteristic feature of DENV is the clear distinction between infection and disease. Although serologic studies have shown high seroprevalence rates worldwide, reaching up to 100% in tropical, DENV-endemic areas, only a small fraction of DENV-infected patients develop symptomatic disease^3^. Even in patients who develop clinical symptoms of dengue disease, only 10-15% progress to severe hemorrhagic disease, suggesting the presence of host risk factors for disease development and severity. The currently accepted model supports the notion that pre-existing anti-DENV IgG antibodies promote the uptake and infection of FcγR-expressing cells to increase susceptibility to dengue disease during secondary DENV infection. However, this model cannot explain why symptomatic dengue disease exhibits such diverse spectrum of clinical phenotypes and only a small fraction of symptomatic patients progress to severe disease. Studies supporting this model have primarily relied on *in vitro* experimental assays based on cell lines that often express a limited set of FcγRs, and their only read-out is the capacity of anti-DENV antibodies to promote infection *in vitro*. Such experimental systems do not take into account the diversity of FcγR-expressing effector cells present *in vivo*, the heterogeneity of the IgG Fc domain and its capacity to interact with the different classes of FcγRs with variable affinity, as well as the as the wide spectrum of effector functions mediated through Fc-FcγR interactions.

To address this limitation, we developed unique mouse models of dengue disease pathogenesis. Such models are based on strains that express the full array of human FcγRs, thereby overcoming long-standing translational barriers associated with the significant interspecies differences between human and mouse FcγR structure and function. To support productive DENV replication, these strains are defective in type I interferon signaling (*Ifnar1*^*-/-*^), which has been previously shown to be sufficient for overcoming the inherent resistance of wild-type mice for DENV infection^34^. In contrast to the AG129 strain, which lacks both IFN-α/β and IFN-γ receptors and has been historically used as the model of choice in studies on the mechanisms of ADE of dengue disease, DENV infection of *Ifnar1*^-/-^ mice induces a disease phenotype that more closely resembles human dengue disease, featuring multiple clinical and pathological manifestations, such as severe thrombocytopenia and hepatocellular necrosis, with minimal involvement of the central nervous system^35^. Also, while IFNγ is known to regulate FcγR expression, and consequently is dysregulated in the AG129 strain, type I IFN signaling has no impact on FcγR expression and therefore *Ifnar1*^-/-^ strains exhibit physiological FcγR expression^17,36^.

Through a combination of human Fc domain variants with defined affinity for specific human FcγR classes, as well as *in vitro* and *in vivo* experimental systems featuring the full array of human FcγRs, we evaluated the role of human FcγRs during the ADE of dengue infection and disease. In contrast to previous reports^29,37-39^, our studies revealed a key role for FcγRIII in promoting increased DENV infection *in vitro*, as well as enhanced disease *in vivo*. Such discrepancy primarily stems from the limitations of experimental systems that have been used in the past to determine the ADE mechanisms of DENV. For example, in studies using function-blocking mAbs, FcγRIIa has been previously identified as the main receptor that mediates ADE of DENV infection *in vitro*. However, this finding likely reflects the use of FcγRIII-negative cell lines, which ignore any potential role for FcγRIII in this process^37,38^. Likewise, although *in vitro* and *in vivo* studies have previously demonstrated an absolute requirement of Fc-FcγR interactions for ADE of dengue infection and disease, there have been conflicting conclusions about the role of the different FcγR pathways in this process. For example, FcγRIIb has been shown to inhibit ADE of dengue infection *in vitro*^29^, whereas other reports suggested that the *in vivo* pathogenic activity of anti-DENV antibodies is the outcome of increased infectivity of liver sinusoidal endothelial cells, which express the inhibitory FcγRIIb^37,39^.

These findings highlight the importance of species-matched experimental systems for the evaluation of human IgG function *in vivo*. Indeed, due to IgG-FcγR species mismatch, conventional mouse strains do not allow the assessment of the ADE activity of polyclonal IgG samples derived from dengue patients. Importantly, the impact of post-translational Fc domain modifications, such as fucosylation, which modulates dengue disease susceptibility, can only be assessed in the context of human FcγRs, as these modifications specifically influence the IgG affinity for human FcγRs^40^. Conversely, the use of mouse mAbs to dissect the ADE mechanisms *in vivo* has limited biological relevance, as differences in the Fc effector function among mouse IgG subclasses are specific for mouse, but not human FcγRs.

In summary, over the past few years, studies on the Fc domain heterogeneity of IgG antibodies that are elicited in dengue patients revisited the concept of ADE and provided evidence to support the association of specific Fc glycan modifications with dengue disease susceptibility. Indeed, the fucosylation status of the Fc domain has been identified as a key determinant of susceptibility to symptomatic dengue disease, highlighting the importance of FcγR pathways in modulating dengue disease pathogenesis. Despite the association of afucosylated anti-DENV IgG antibodies with dengue disease severity, clear evidence on their *in vivo* pathogenic activity has been missing, due to the paucity of appropriate animal models that could accurately evaluate human FcγR function. Through the development and use of mouse strains with high translational relevance to human FcγR biology, our studies revealed a major role for FcγRIIIa, which through its capacity to interact with higher affinity with afucosylated Fc glycoforms modulates dengue disease pathogenesis. These findings highlight the importance of the FcγRIIIa-afucosylated Fc axis in dengue disease, having important implications for the development of strategies to mitigate the risk for ADE among vaccinated individuals, as well as to prevent or control symptomatic disease in high-risk populations.

## Acknowledgments

We thank P. Smith, E. Lam, R. Peraza, R. Francis, and J. Edgar for technical assistance, all the members of the Laboratory of Molecular Genetics and Immunology (Rockefeller University) for discussions, S. Carrasco, S. St. Jean, and staff from the Laboratory Comparative Pathology for histopathology support (with funding from NIH Core Grant P30CA008748), M. Mack (University Hospital Regensburg) for providing us the anti-CCR2 antibody (clone MC-21), the CRISPR & Genome Editing Center at Rockefeller University for their help with the design of the *Ifnar1*^-/-^ KO mice, and The Rockefeller University for continued institutional support. The following reagent was obtained through BEI Resources, NIAID, NIH: Dengue Virus Type 2 (DENV2), New Guinea C (NGC), NR-84. Research reported in this publication was supported by the National Institute of Allergy and Infectious Diseases Grants R01AI137276 (to S.B.), U19AI111825 (to J.V.R., C.M.R, and S.B.), and R01AI124690 (to C.M.R.). The content is solely the responsibility of the authors and does not necessarily represent the official views of the NIH.

## Author contributions

Conceptualization: R.Y., J.V.R., and S.B.; Methodology: R.Y. and K.S.K.; Investigation and Resources: R.Y. and K.S.K.; Writing: R.Y., J.V.R., and S.B.; Visualization: R.Y. and S.B.; Intellectual input: M.R.M. and C.M.R.; Supervision: J.V.R. and S.B.; Funding Acquisition: J.V.R. and S.B.

## Extended Data

**Extended Data Fig. 1:**
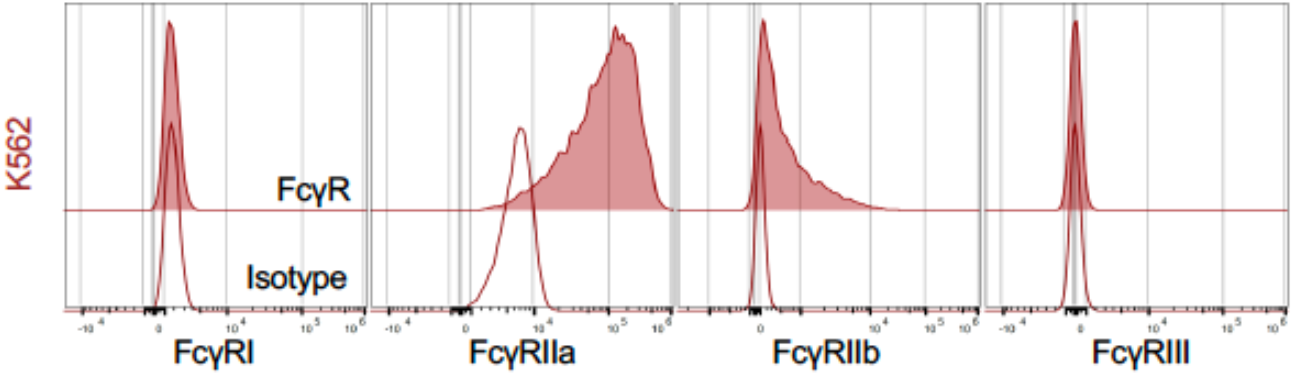
FcγR expression profile of K562 cells. FcγR expression on K562 was assessed by flow cytometry using antibodies against FcγRI, FcγRIIa, FcγRIIb, and FcγRIII (shaded histogram). Corresponding isotype control is shown in open histograms.

**Extended Data Fig. 2:**
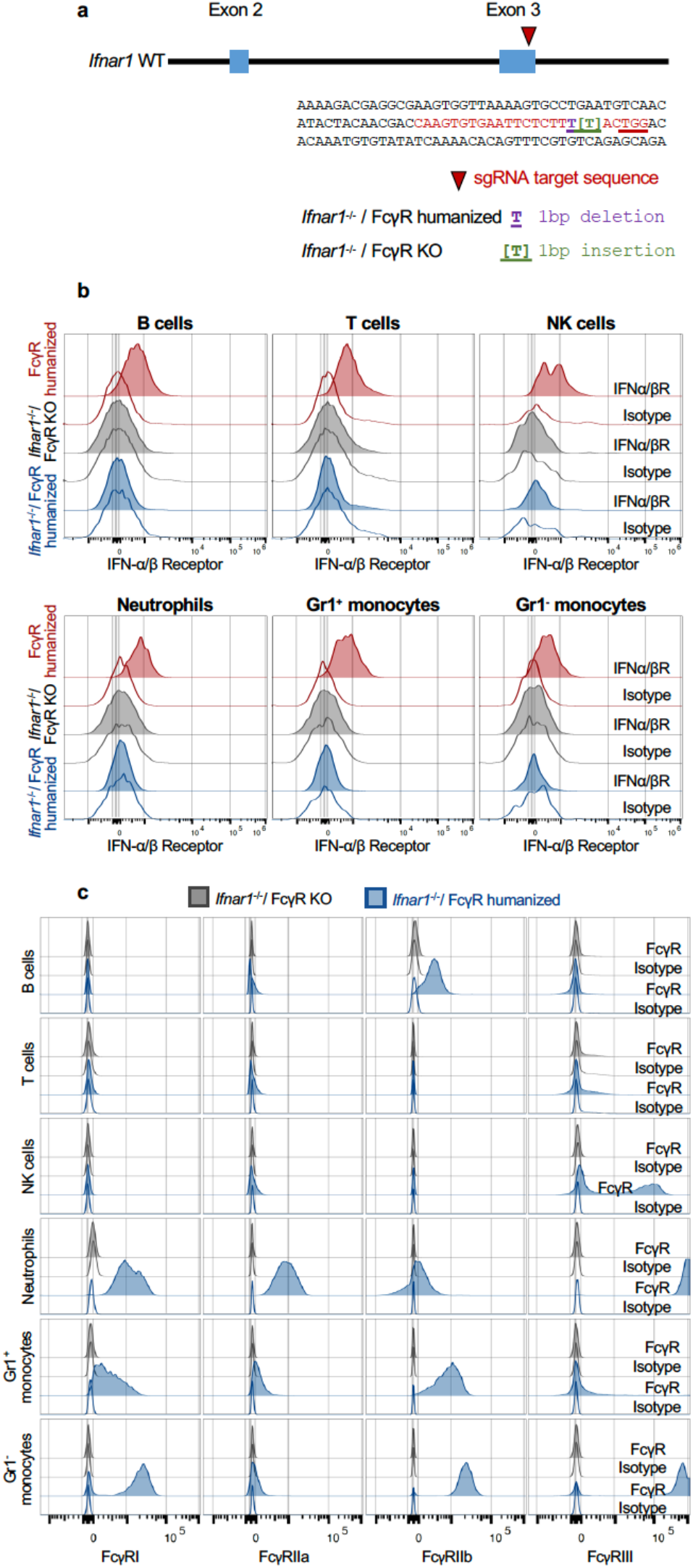
Characterization of *Ifnar1*^-/-^/FcγR KO and *Ifnar1*^-/-^/ humanized mice. (**a**) Overview of *Ifnar1* gene targeting with CRISPR/Cas9 in mice. Single guide RNA (sgRNA) was designed to target a sequence in exon 3 of the mouse *Ifnar1*ifnar1 gene (marked in red). CRISPR-Cas9 constructs were injected to FcγR KO mouse or FcγR humanized mouse and resulted in 1 bp insertion or 1 bp deletion, respectively, that led to a premature stop codon. (**b**) IFNα/βR expression was assessed by flow cytometry in FcγR KO (gray) or FcγR humanized (blue) mice in comparison to FcγR humanized mice (red). (**c**) FcγR expression in various leukocyte populations in the blood was assessed by flow cytometry for *Ifnar1*^-/-^/FcγR KO mice (gray) and *Ifnar1*^-/-^/FcγR humanized mice (blue). Matching isotype controls are indicated by open histograms.

**Extended Data Table 1:** Affinity of human IgG1 Fc domain variants for the various classes of human FcγRs.

| Human IgG1<br>Fc variant | Activating |  |  |  | Inhibitory |  |
| --- | --- | --- | --- | --- | --- | --- |
|  | FcγRIIa |  | FcγRIIIa |  | FcγRIIb |  |
|  | KD (M) | Fold | KD (M) | Fold | KD (M) | Fold |
| Wild-type | 1.03×10<sup>-6</sup> | 1.0 | 7.4×10<sup>-7</sup> | 1.0 | 3.11×10<sup>-6</sup> | 1.0 |
| GRLR | n.d.b. |  | n.d.b. |  | n.d.b. |  |
| V11 | 1.4×10<sup>-6</sup> | 0.7 | 7.15×10<sup>-7</sup> | 1.0 | **1.12×10<sup>-7</sup>** | **27.8** |
| GAALIE | **3.87×10<sup>-7</sup>** | **2.7** | **2.18×10<sup>-7</sup>** | **3.4** | 4.15×10<sup>-6</sup> | 0.8 |
| GA | **1.86×10<sup>-7</sup>** | **5.5** | 9.03×10<sup>-7</sup> | 0.8 | 2.89×10<sup>-6</sup> | 1.1 |
| ALIE | 2.07×10<sup>-6</sup> | 0.5 | **1.69×10<sup>-7</sup>** | **4.4** | 3.92×10<sup>-6</sup> | 0.8 |
| Afucosylated | 7.47×10<sup>-7</sup> | 1.4 | **1.45×10<sup>-7</sup>** | **5.1** | 2.22×10<sup>-6</sup> | 1.4 |

n.d.b.: no detectable binding

**Extended Data Fig. 3:**
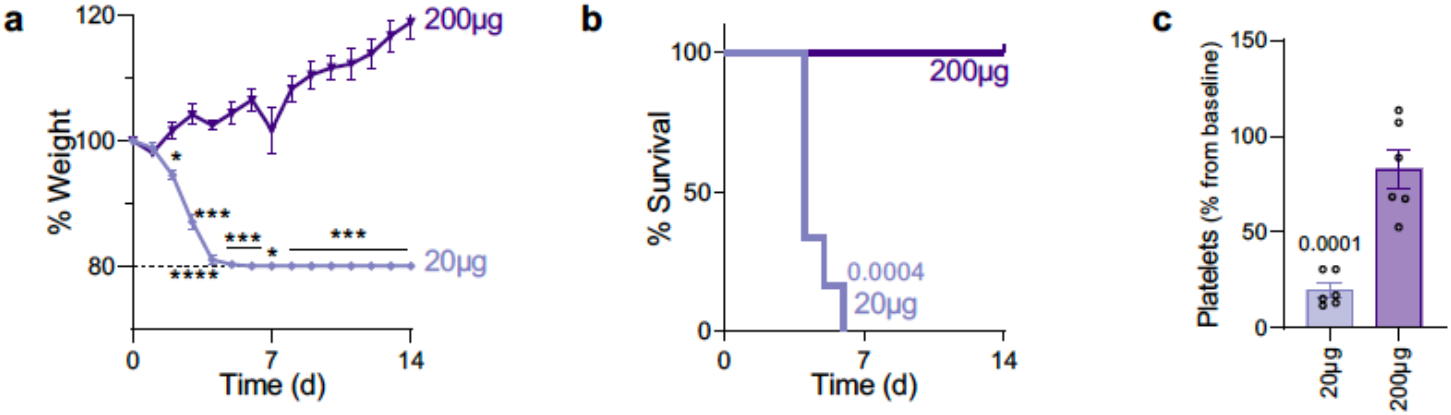
Titration of anti-DENV 2D22 mAb in vivo. (**a-c**) *Ifnar1*^-/-^/FcγR humanized mice were administered i.v. with 20 μg or 200 μg of anti-DENV 2D22 mAb 8h prior to DENV2 challenge. Weight loss (**a**), survival (**b**), and platelet counts (**c**) were compared between the two doses by two-way ANOVA (Bonferroni post hoc analysis adjusted for multiple comparisons), log-rank (Mantel-Cox) test and t-test, respectively. \**P*<0.03; \*\*\**P*<0.0008; \*\*\*\**P*<0.0001.

## Methods

### Viruses and cell lines

K562 cells (ATCC, CRL-3343), U937 cells (ATCC, CRL-1593.2) and U937-DC-SIGN (ATCC, CRL-3253) cells were cultured in Roswell Park Memorial Institute (RPMI) 1640 Medium supplemented with 10% FBS, 50 U/ml penicillin and 50 μg/ml streptomycin (ThermoFisher). U937-panFcγR cells were cultured in RPMI 1640 supplemented with 10% FBS, 50 U/ml penicillin, 50 μg/ml streptomycin and 5 mg/ml blasticidin (InvivoGen). All cell lines were maintained at 37°C at 5% CO_2_ incubator. C6/36 cells (ATCC, CRL-1660) were cultured in Minimum Essential Medium (MEM) α medium supplemented with 10% FBS, 50 U/ml penicillin and 50 μg/ml streptomycin (ThermoFisher) and maintained at 29°C at 5% CO_2_ incubator. Expi293F cells (ThermoFisher) were maintained at 37°C, 8% CO_2_ in Expi293 expression medium (ThermoFisher) supplemented with 10 U/ml penicillin and 10 μg/ml streptomycin. All cell lines tested negative for mycoplasma contamination.

DENV2 New Guinea C (BEI Resources, NR-84) was amplified in C6/36 cells at 29°C for 5 d. The virus-containing supernatant was harvested, clarified by centrifugation (3,000*g*; 10 min), filtered using a disposable vacuum filter system with a 0.45 μm membrane, and then concentrated ten-fold using Amicon Ultra 100K. RNA levels were determined by TaqMan Fast Virus 1-Step Master Mix with specifically designed primers and a TaqMan probe that bind a conserved region in the envelope (E) gene of DENV2 (Forward primer: 5’-CAGGCTATGGCACYGTCACGAT-3’; Reverse primer: 5’-CCATYTGCAGCARCACCATCTC-3’; Probe: 5’-FAM/ZEN/CTCYCCRAGAACGGGCCTCGACTTCAA/IBFQ-3’)^41^. Quantitative PCR was performed using an Applied Biosystems QuantStudio 6 Flex cycler using the following parameters: 50°C for 5 min, 95°C for 20 s, followed by 45 cycles of 95°C for 3 s, and 30 s at 60°C. To calculate viral titer, an RNA standard curve was generated from a defined viral stock (ATCC, VR-3229SD) and titer was expressed as genome equivalent (GE).

### Animals

Animal care and experimentation were consistent with federal laws and were approved by the Rockefeller University Institutional Animal Care and Use Committee. Mice were maintained at the Comparative Bioscience Center at the Rockefeller University at a controlled ambient temperature (20-25°C) and humidity (30–70%) environment with 12-h dark:light cycle. *Ifnar1*^-/-^ mice were generated in the FcγR knockout (mFcγRα^-/-^; *Fcgr1*^-/-^) and FcγR humanized mice (mFcγRα^-/-^, *Fcgr1*^-/-^, hFcγRI^+^, hFcγRIIa_R131_+, hFcγRIIb^+^, hFcγRIIIa_F158_^+^ and hFcγRIIIb^+^) background. Guide RNA (CAAGTGTGAATTCTCTTTACTGG) was designed to target a sequence in exon 3 of the *Ifnar1* gene. This guide was subsequently delivered as a RNP complex (crRNA+tracrRNA+Cas9 protein) into mouse zygote via zygote injection. Target DNA from F0 mice was amplified and cloned into TOPO-TA vector and sequenced by Sanger sequencing. A 1 bp deletion was found in the *Ifnar1*^-/-^/FcγR humanized mice while 1 bp insertion was found in *Ifnar1*^-/-^/FcγR KO mice. Individual mutant founders were crossed to FcγR humanized or FcγR KO mice, respectively. Following the initial screening, TaqMan SNP Genotyping Assay was designed for each indel mutation (assay ID AN9HN3G for *Ifnar1*^-/-^/FcγR humanized mice and ANRWJMM for *Ifnar1*^-/-^/FcγR KO mice) and mice were genotyped by TaqMan Genotyping Master Mix (ThermoFisher).

### Generation of recombinant antibodies

Site-directed mutagenesis (SDM) using specific primers was used to generate the various human IgG1 Fc variants^26^. Afucosylated human IgG1 was generated in the presence of 0.2 mM 2-deoxy-2-fluoro-l-fucose (2FF) (Carbosynth, MD06089) during transfection of recombinant antibodies^42^. Recombinant IgG antibodies were produced in Expi293 cells as previously described^43^. Expi293 were transfected with 1:1 ratio of heavy chain:light chain constructs using ExpiFectamine 293 transfection kit. Seven days post-transfection, the supernatant was collected, centrifuged to remove cell debris, and sterile filtered through 0.22 μm filter. Antibodies were purified on protein G affinity purification column and purity was assessed by SDS-PAGE followed by SafeStain blue staining (ThermoFisher). All antibody preparations were >95% pure.

### DENV neutralization assay

Neutralization activity of IgG1 Fc variants of anti-DENV mAbs was measured in U937-DC-SIGN cells^44^. Antibodies were serially diluted three-fold starting at 10 μg/ml, and pre-incubated with DENV (1.75×10^7^ GE/well) at 37°C for 20 min in 96 U-shaped plates. U937-DC-SIGN cells were added to each well at a concentration of 2×10^4^ cells/well and then incubated overnight at 37°C. One day post-infection cells were centrifuged (500*g*, 5 min), carefully washed with PBS and then fixed using Cytofix/Cytoperm buffer (BD). Cells were then stained with DyLight 650-conjugated 4G2 (ATCC, HB-112) and analyzed by flow cytometry. The antibody dilution that neutralized 50% of the viruses was calculated by nonlinear, dose-response regression analysis.

### *In vitro* ADE assay

ADE of DENV infection was measured in K562, U937, or U937-panFcγR cells. Antibodies were serially diluted (10-fold starting at 10 μg/ml) and pre-incubated with DENV (3.5×10^7^ GE/well) at 37°C for 20 min in 96U-shaped plates. Tested cells were added to each well at a concentration of 10^4^ cells/well and then incubated overnight at 37°C. One day post-infection, cells were centrifuged (500*g*, 5 min), carefully washed with PBS and then fixed using Cytofix/Cytoperm buffer. Cells were stained by flow cytometry with DyLight 650-conjugated 4G2. AUC was calculated and one-way ANOVA (Bonferroni post hoc analysis adjusted for multiple comparisons) was used for statistical analysis.

### Flow cytometry

K562, U937, and U937-panFcγR cells were washed once with PBS and stained with the following antibodies (all used at 5 μg/ml): anti-human FcγRI (clone 10.1)-BrilliantViolet605, anti-human FcγRIIa (clone IV.3)-DyLight488, anti-human FcγRIIb (clone 2B6)-Dylight650 and anti-human FcγRIIIa/b (clone 3G8)-BrilliantViolet650.

Red blood cells (RBC) were lysed (RBC lysis buffer, Biolegend) and resuspended in staining buffer (PBS containing 0.5% (w/v) BSA and 5 mM EDTA). Cells were then labeled with the following antibodies (1:100 unless otherwise stated): anti-B220-PE-Cy5, anti-Gr1-BrilliantViolet421, anti-CD3-BrilliantViolet650, anti-CD11b-BrilliantViolet711, anti-NK1.1–PE– Cy7, anti-human FcγRI (clone 10.1)-BrilliantViolet605 (used at 5 μg/ml), anti-human FcγRIIa (clone IV.3)-DyLight488 (5 μg/ml), anti-human FcγRIIb (clone 2B6)-Dylight650 (5 μg/ml) and anti-human FcγRIIIa/b (clone 3G8)-PE (5 μg/ml).

Spleens were harvested on day 4 post-infection and homogenized by mechanical shearing. Following RBC lysis, single cell suspensions were labeled with LIVE/DEAD Fixable Near-IR (ThermoFisher) and resuspended in PBS containing 0.5% (w/v) BSA and 5 mM EDTA. Cells were labelled with the following mixture of fluorescently labelled antibodies (1:200 unless otherwise stated): anti-Siglec F-PerCP-eFluor710, anti-CD11b-AlexaFluor700, anti-F4/80-BrilliantViolet711, anti-CD3-BrilliantViolet510, anti-CD19-BrilliantViolet510, anti-NK1.1-BrilliantViolet510, and anti-Ly6G-BrilliantViolet510. For the evaluation of dengue infection DyLight650-conjugated 2H2 (ATCC, HB-114) was added to the mixture (used at 10 μg/ml). for the characterization of FcγR expression the following antibodies were added to the mixture: anti-human FcγRI (clone 10.1)-BrilliantViolet605 (used at 5 μg/ml), anti-human FcγRIIa (clone IV.3)-DyLight488 (5 μg/ml), anti-human FcγRIIb (clone 2B6)-Dylight650 (5 μg/ml) and anti-human FcγRIIIa/b (clone 3G8)-BrilliantViolet650 (5 μg/ml).

The following isotype control antibodies were used for both panels: mouse IgG1 isotype control-Dylight650 (used at 5 μg/ml), mouse IgG2b kappa isotype control–DyLight488, mouse IgG1 kappa isotype control–PE (or mouse IgG1 kappa isotype control–BrilliantViolet650), mouse IgG1 kappa isotype control–BrilliantViolet605. Unlabeled anti-FcγR blocking antibodies were added to samples that were stained with isotype control antibodies as follow: anti-human FcγRI (clone 10.1), anti-human FcγRIIa (clone IV.3), anti-human FcγRIIb (clone 2B6), and anti-human FcγRIIIa/b (clone 3G8) (used at 5 μg/ml and incubated for 5 min before staining with fluorescently labelled antibodies). Samples were analyzed on an Attune NxT flow cytometer (ThermoFisher) using Attune NxT software v3.1.2 and data were analyzed using FlowJo (v10.8.1) software.

### *In vivo* DENV challenge

Animal infection experiments were performed at the Comparative Bioscience Center of the Rockefeller University in animal biosafety level 2 (ABSL-2) containment in compliance with institutional and federal guidelines. Mice (males and females; 4-5 weeks old) were anaesthetized with isoflurane (3%) in a VetFlo high-flow vaporizer before challenge with DENV (New Guinea C strain, 3.5×10^8^ GE, i.v.). For *in vivo* ADE assays, antibodies (monoclonal anti-DENV or polyclonal IgG antibodies purified from hospitalized dengue patients (n=35) as described previously^19^) were administered i.v. 8h before virus challenge. After infection, mice were monitored daily, and their weights were recorded for up to 14 days. Death was determined by a 20% body weight loss threshold that was authorized by the Rockefeller University Institutional Animal Care and Use Committee. Before treatment, mice were randomized based on sex.

### Determination of tissue viral titers

Mice were euthanized on day 4 post-infection and spleens were harvested. Tissue was lysed in Trizol (ThermoFisher) and dissociated in gentle MACS M tubes using the gentleMACS Octo Dissociator (Miltenyi Biotec). Samples were transferred into Phasemaker tubes (ThermoFisher) and chloroform was added (200 μl chloroform per ml TRIzol). After vigorous shaking, tubes were rested for 5 min and then centrifuged for 15 min at 12,000*g* at 4°C. The aqueous phase containing the RNA was transferred into a new tube and RNA extraction was performed by using RNeasy 96 kit (Qiagen). DENV titers were determined by quantitative PCR with reverse transcription assay using TaqMan Fast Virus 1-Step Master Mix, and the same primers-probe mixture that was used for virus quantification (described in “Viruses and cell lines” section). Quantitative PCR was performed using an Applied Biosystems QuantStudio 6 Flex cycler using the following parameters: 50°C for 5 min, 95°C for 20 s, followed by 45 cycles of 95°C for 3 s, and 30 s at 60°C. Signal from unknown samples was compared to a known RNA standard curve (ATCC, VR-3229SD) and viral titers were expressed as GE per mg tissue.

### Measurement of serum cytokine levels

Blood was collected from the retro-orbital sinus of mice under isoflurane anesthesia and placed into gel microvette tubes. Serum was fractionated by centrifugation (10,000*g*, 5 min) and stored at -20°C. The levels of cytokines in the serum samples were determined by Cytokine & Chemokine Convenience 36-Plex Mouse ProcartaPlex™ Panel 1A (ThermoFisher) according to the manufacturer’s instructions.

### *In vivo* depletion

NK cells, neutrophils, or CCR2^+^ monocytes were depleted in mice by administration of anti-NK1.1 (clone PK136, 150 μg), anti Gr-1 (clone RB6-8C5, 150 μg), or anti-CCR2 (clone MC-21, 50 μg) mAbs, respectively. Macrophages were depleted by administration of 100 μl clodronate liposomes. All depletion reagents were administered 24 hours before DENV challenge, i.v.

### Splenectomy

Mice were anaesthetized with a ketamine (75 mg/kg) and xylazine (15 mg/kg) mixture (administered intraperitoneally) before spleen removal. A longitudinal incision was made on the left dorsolateral side of the abdomen. The splenic arteries were cauterized, and the spleen was removed. Sham surgery was performed by exposing the spleen and then closing the abdominal cavity. The skin was sutured following closure of the abdominal wall^45^. After the surgery, the animals were allowed to recover for 5 days prior to antibody administration and DENV challenge.

### Surface plasmon resonance

All SPR experiments were performed with a Biacore T200 SPR system (GE Healthcare) at 25°C in HBS-EP+ buffer (10 mM Hepes, pH 7.4, 150 mM NaCl, 3.4 mM EDTA, 0.005% [v/v] surfactant P20). Monoclonal antibodies were immobilized on Series S Protein G sensor chip (GE Healthcare) at a density of 2,000 response units (RU). Serial dilutions of soluble human FcγR ectodomains were injected through the flow cells at a flow rate of 20 μl/min with the concentration ranging from 1,024 to 8 nM (serial two-fold dilutions). Association time was 60 s followed by a 600-s dissociation. At the end of each cycle, the sensor surface was regenerated with glycine HCl buffer (10 mM, pH 1.5; 50 μl/min for 30 s). Background binding to blank immobilized flow cells was subtracted, and the steady-state affinity model was used to validate that the equilibrium had been reached under the selected concentration series of the analytes (RUmax vs. analyte concentration). Affinity constants (KD) were then calculated using BIAcore T200 evaluation software v.2.0 (GE Healthcare).

### Hematological analysis

Approximately 100 μl blood samples were collected from the retro-orbital sinus of mice under isoflurane anesthesia. Blood was placed in BD Microtainer tubes (coated with ethylenediaminetetraacetic acid (K2-EDTA) and platelet counts were measured by Element HT5 hematology analyzer (Heska).

### Histopathological analysis

Organs (liver, spleen, lung, kidney, intestine) from euthanized mice were fixed for 5 days by submersion in 10% formalin. Fixed tissues were embedded in paraffin, sectioned at 4-μm thickness, and stained with haematoxylin and eosin. Sections from different organs were microscopically evaluated by a board-certified veterinary anatomic pathologist and representative images were captured with an Olympus BX45 light microscope using an SC30 camera with the cellSens Dimension software.

### Statistical analysis

Results from multiple animals or experiments are presented as mean ± s.e.m. One-or two-way ANOVA was used to test for differences in the mean values of quantitative variables, and where statistically significant effects were found, post hoc analysis using Bonferroni (adjusted for multiple comparisons) test was performed. Two-tailed t-test was used to test for differences in datasets with two groups. Statistical differences between survival rates were analyzed by comparing Kaplan–Meier curves using the log-rank (Mantel–Cox) test. Data were collected and analyzed with Microsoft Excel and GraphPad Prism v.9.4.1 software (GraphPad) and P < 0.05 was considered to be statistically significant.

